## Supplementary for "Heterologous HSPC transplantation rescues neuroinflammation and ameliorates peripheral manifestations in the mouse model of lysosomal transmembrane enzyme deficiency, MPS IIIC"

### Supplementary materials

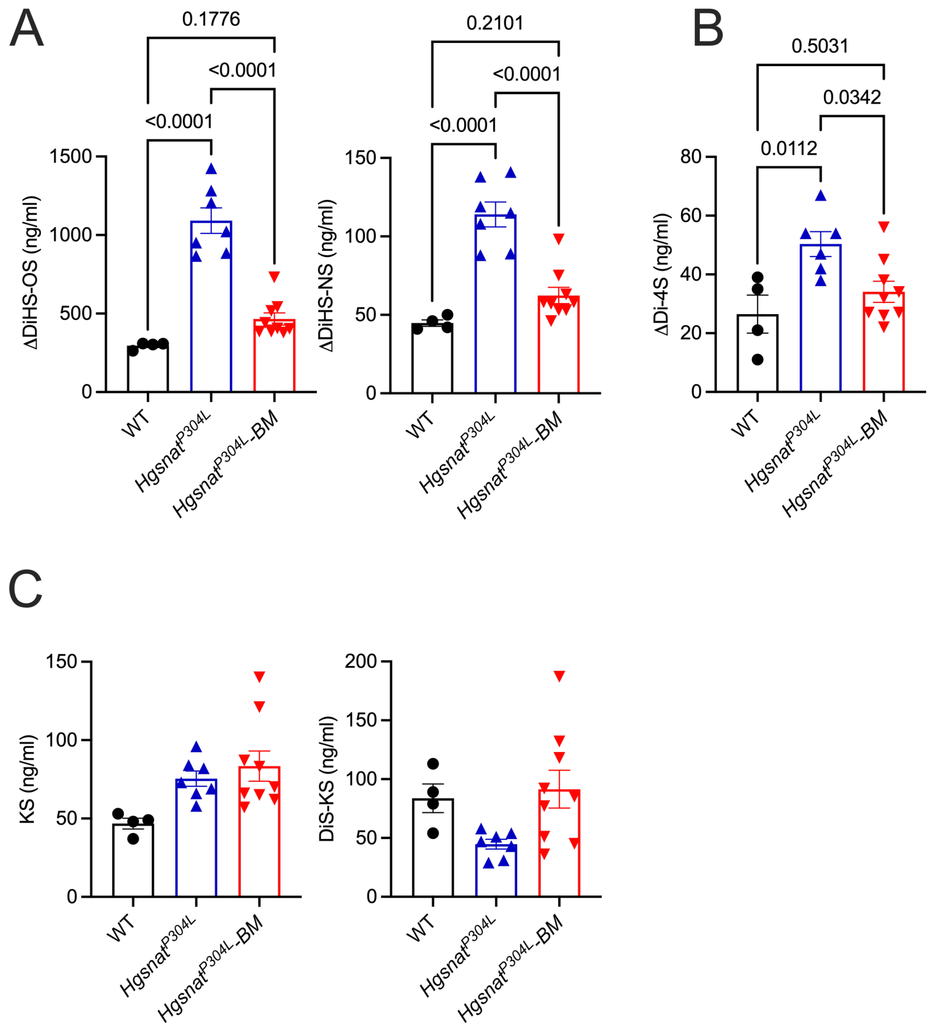


**Figure S1. Levels of disaccharides produced by enzymatic digestion of HS, KS and DS in dry blood spots of mice 6 weeks after HSPC transplantation.**

Levels of HS-derived O-sulfated (ΔDiHS-OS) and N-sulfated (ΔDiHS-NS) disaccharides **(A)** or dermatan sulfate-derived disaccharide (ΔDi-4S) **(B)** are increased in the DBS of untreated *Hgsnat$^{P304L}$* mice, while in the transplanted *Hgsnat$^{P304L}$* mice of the same age, the levels of all three disaccharides are not significantly different from the normal levels. Levels of disaccharides produced by enzymatic digestion of mono (KS) and di-sulfated (DiS-KS) keratan sulfate are similar in DBS of WT, *Hgsnat$^{P304L}$* and transplanted *Hgsnat$^{P304L}$* mice. All graphs show individual data, means and SD of experiments performed with samples of 4-9 male and female mice per genotype per treatment. P values were calculated by one-way ANOVA with Tukey post hoc test.

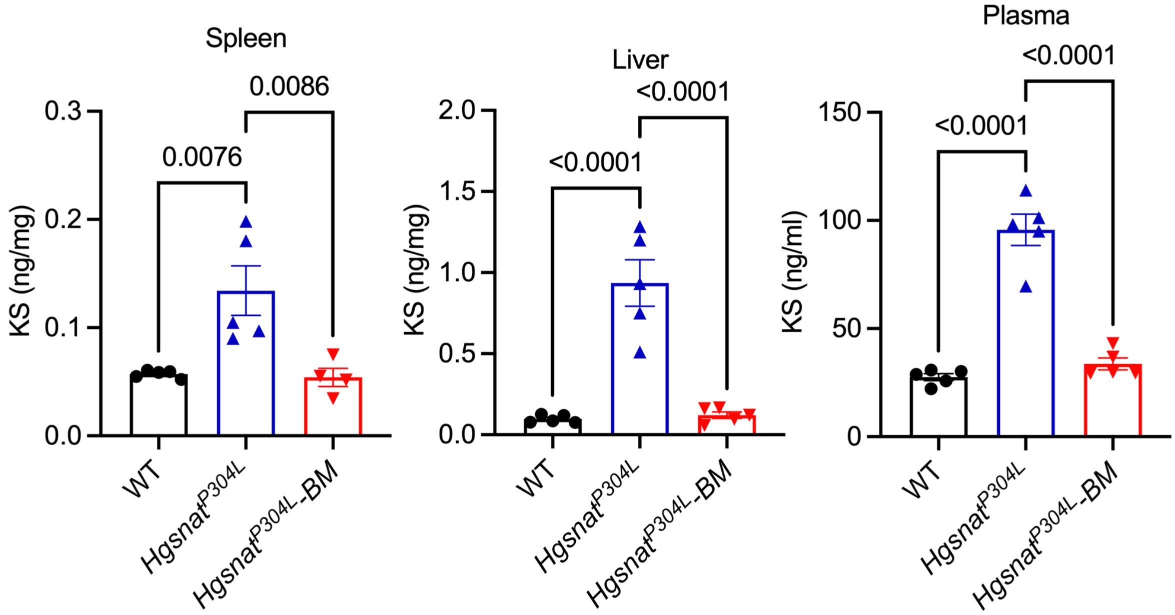


**Figure S2. Levels of disaccharides produced by enzymatic digestion of mono-sulfated KS in blood plasma, liver and spleen of WT, *Hgsnat$^{P304L}$* and transplanted *Hgsnat$^{P304L}$* mice at the age of 8 months.**

All graphs show individual data, means and SD of experiments performed using tissues from 5 male and female mice per genotype per treatment. P values were calculated by one-way ANOVA with Tukey post hoc test.

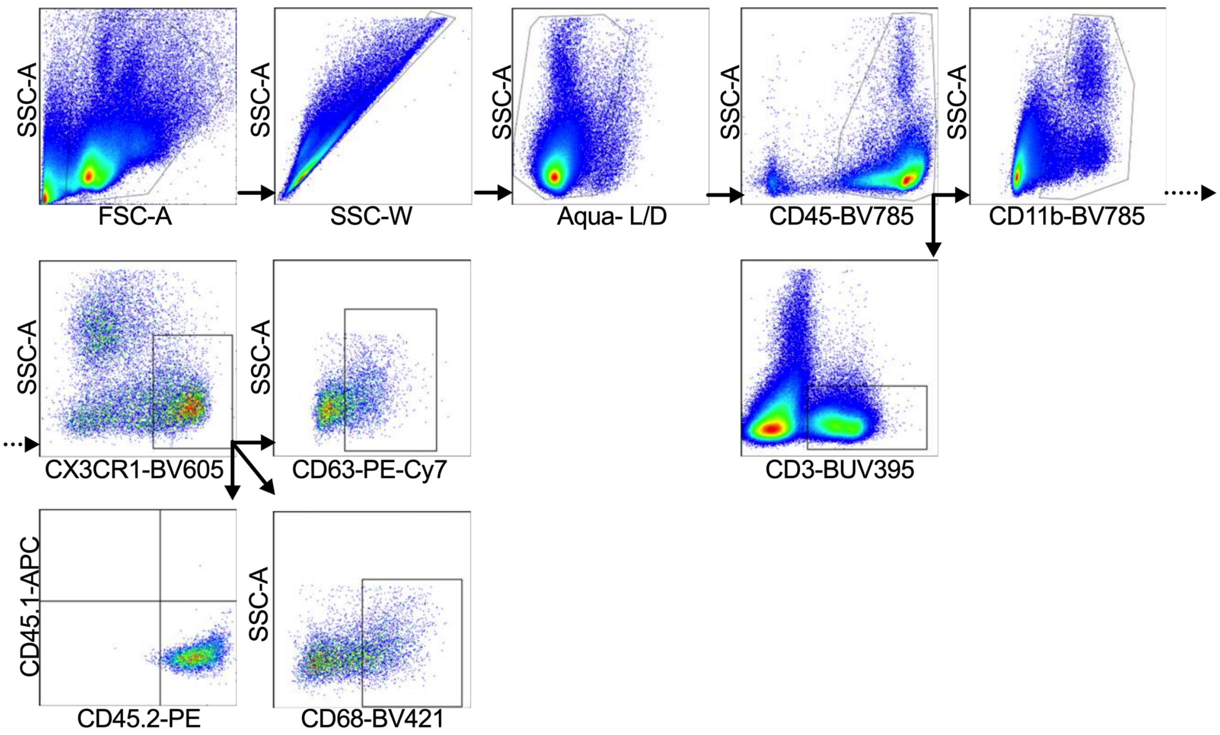


**Figure S3. Gating strategy for the analysis of dissociated brain/spinal cord cells and splenocytes by flow cytometry.**

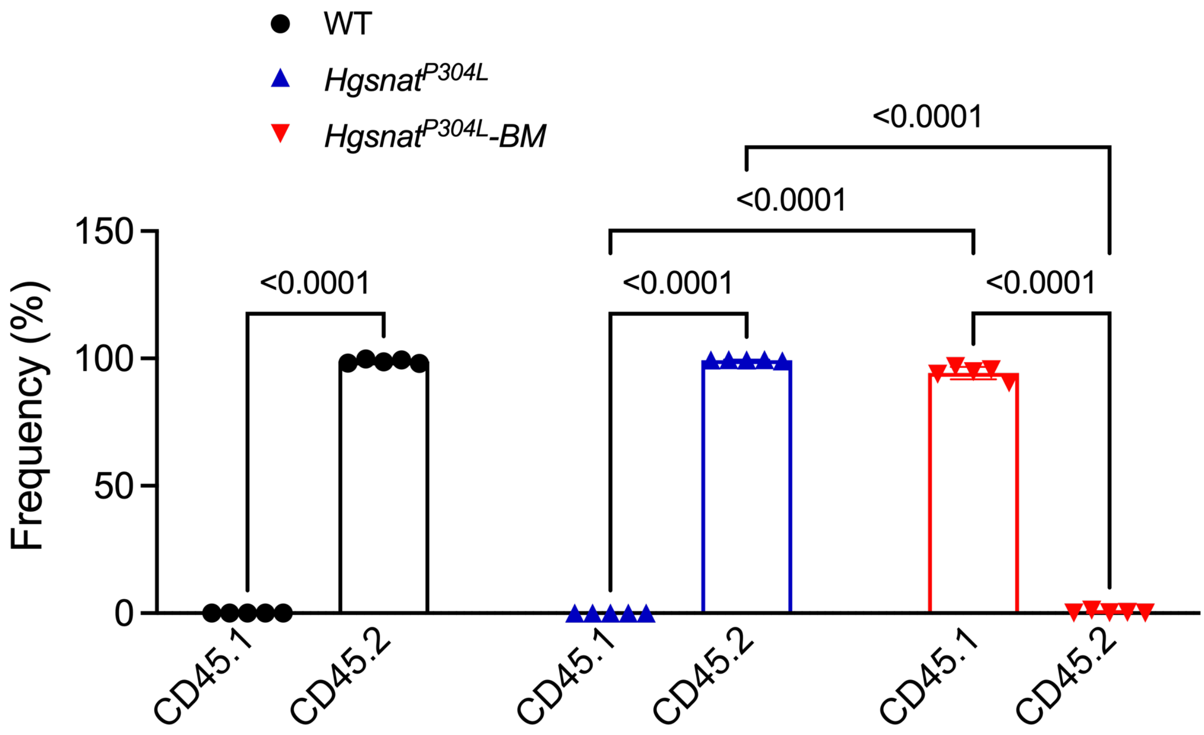


**Figure S4. All CD45/CD11b/CX3CR1-positive macrophages in the spleen of transplanted *Hgsnat*$^{P304L}$ mice show CD45.1 phenotype indicating that they are derived from transplanted HSPC.**

Graphs show individual results, means and SD from experiments conducted with 5 mice per group. P values were calculated using one-way ANOVA test with Tukey post hoc test.

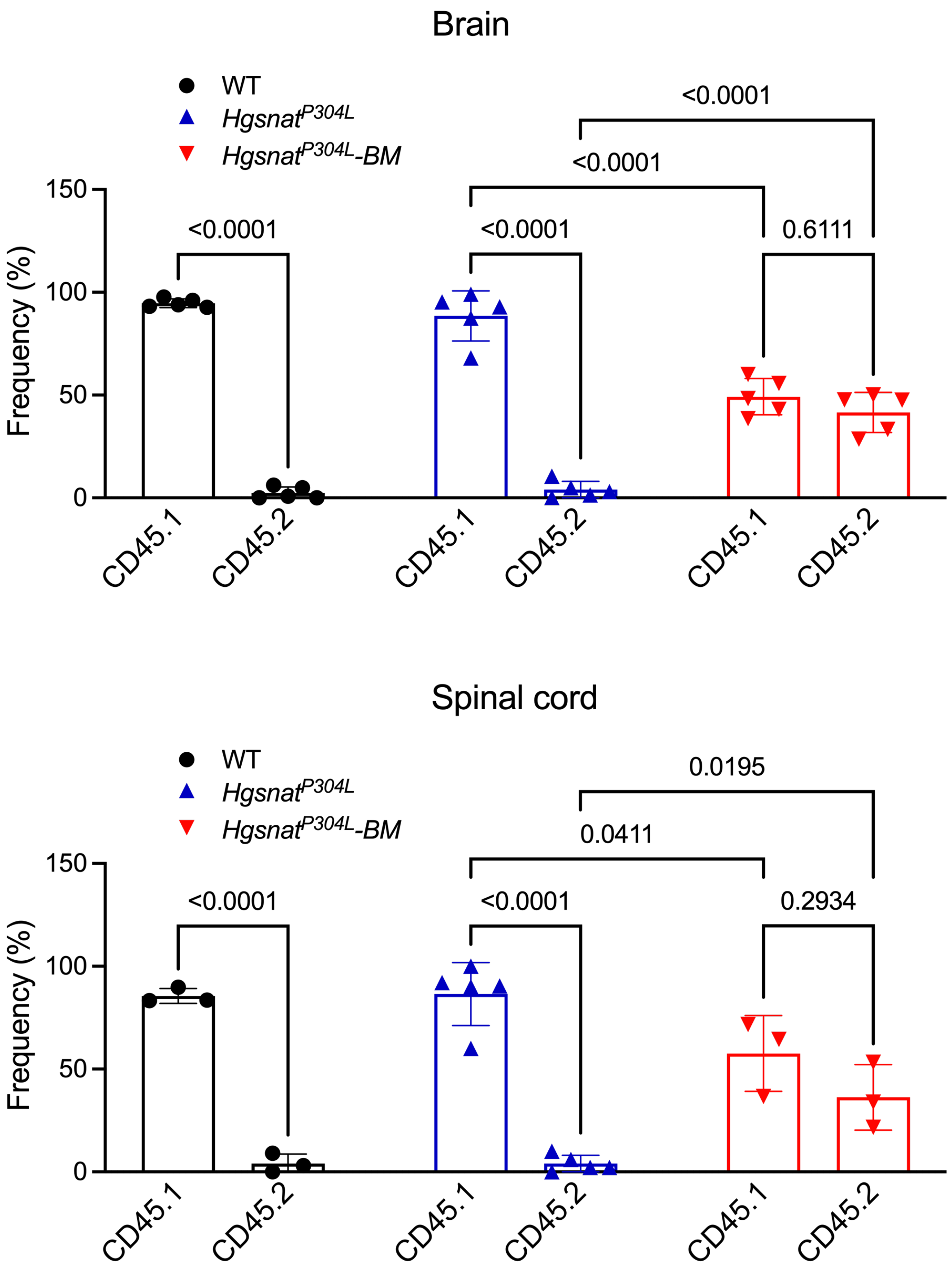


**Figure S5. In both brain and spinal cord tissues approximately 50% of CD45/CD11b/CX3CR1-positive macrophages/microglia cells are CD45.1-positive and derived from transplanted HSPC.**

Graphs show individual results, means and SD from experiments conducted with 3-5 mice per group. P values were calculated using one-way ANOVA test with Tukey post hoc test.

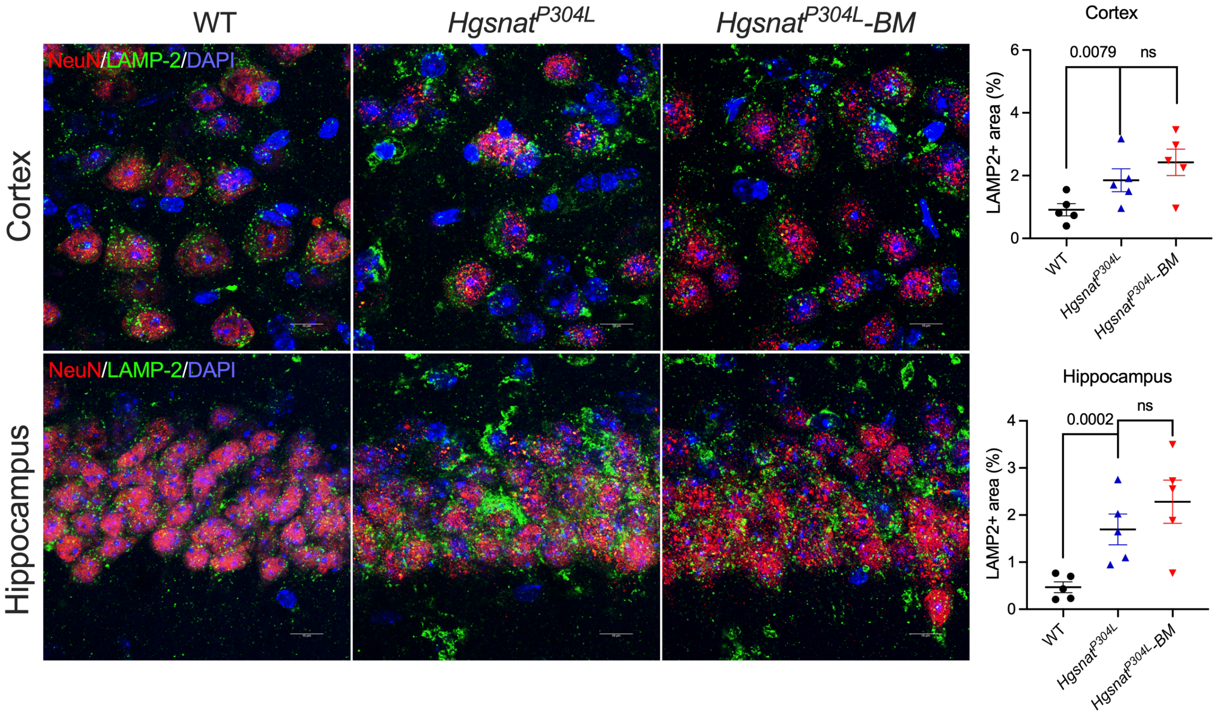


**Fig. S6. Levels of LAMP-2-positive puncta are unchanged in cortical and hippocampal neurons of transplanted *Hgsnat*$^{P304L}$ mice.**

Panels show representative images of brain cortex (layers 4-5) and CA1 region of hippocampus of 8-month-old WT and treated or untreated *Hgsnat*$^{P304L}$ mice labeled for LAMP-2 (green) and NeuN (red). DAPI was used as a nuclear counterstain. Scale bars equal 10 µm. Graphs show quantification of LAMP-2-positive areas in NeuN-positive cells with ImageJ software. All graphs show individual results, means and SD from experiments conducted with 5 mice (three panels per mouse) per genotype per treatment. P values were calculated using Nested one-way ANOVA test with Tukey post hoc test.

**Supplementary Table 1. Mouse engraftment 6 weeks after transplantation**

| Body Weight before transplantation (g) | ID # /sex | CD45.1+ myeloid cells (%) | CD45.2+ myeloid cells (%) |
|---|---|---|---|
| 26 | 11354 / F | 83.7 | 15.1 |
| 30 | 11335 / M | 84.5 | 14.2 |
| 30 | 11334 / M | 88.2 | 10.6 |
| 33 | 11333 / M | 82.0 | 16.8 |
| 23 | 11560 / F | 88.0 | 9.9 |
| 24 | 11561 / F | 87.6 | 12.3 |
| 24 | 11562 / F | 86.8 | 12.3 |
| 30 | 11552 / M* | NA | NA |
| 29 | 11553 / M | 88.3 | 10.6 |
| 30 | 11554 / M | 91.6 | 7.6 |
| 34 | 11556 / M** | NA | NA |
| 29 | 11557 / M | 94.1 | 4.9 |
| 24 | 11566 / F | 90.8 | 7.9 |
| 26 | 11565 / F | 91.4 | 7.0 |
| 25 | 11564 / F | 93.2 | 6.4 |
| 24 | 11563 / F | 91.3 | 8.0 |

* Died on the 4$^{th}$ week after transplantation
** Died on the 3$^{d}$ week after transplantation
